# Modality-specific predictive templates in pre-stimulus EEG activity

**DOI:** 10.1101/2023.09.27.559806

**Authors:** Isabelle Hoxha, Sylvain Chevallier, Arnaud Delorme, Michel-Ange Amorim

## Abstract

Perceptual decision-making is a combination of sensory information and prior beliefs. In order to perform actions in a timely fashion, it is necessary to anticipate the timing at which events occur, but also what event is more likely than the other. While EEG signatures of anticipation have been identified it is less clear whether classifiers trained on informative (cued) trials generalize to uncued trials and whether such decoded templates predict trial-by-trial shifts in decision strategy (e.g., drift-rate or starting-point changes in a Diffusion Decision Model). This study aimed to determine whether human participants anticipated a visual or auditory stimulus at the single-trial level in both cued and uncued trials. We found that pre-stimulus brain activity contains information about the expected upcoming stimulus and that this information can be successfully extracted from single-trial brain activity. Behavioral analyses revealed a connection between correct anticipation and shifts in decision strategy, while also validating the classification of uncued trials. Importantly, the classification of uncued trials confirms that expectations build even in the absence of triggers. These findings highlight the presence of single-trial, stimulus-specific neural signatures of anticipation, offering new insights into trial-to-trial variability in decision-making and advancing our understanding of cognitive processes.

**Highlights:**

- Distributed low-frequency pre-stimulus EEG encodes modality-specific expectations.
- Anticipatory neural states are evoked even in the absence of explicit cues.
- Decoded anticipation predict response speed and accuracy.

## Introduction

Consider a situation in which a researcher is pouring coffee while awaiting an important phone call from a funding agency. Visual attention is directed toward the rising coffee level, whereas auditory attention is oriented toward the ringtone. At some point, an action will be required, but only one sensory modality can be prioritized at a time. Effective behavior in such situations requires not only anticipating when an action will be needed, but also which sensory modality should be preferentially prepared.

Anticipation is a fundamental property of brain function. Rather than passively responding to incoming stimuli, the brain continuously generates predictions about future events in order to optimize perception and action (Petro et al., 2019; Poulton, 1950). Within predictive processing frameworks, expectations are thought to bias sensory systems in advance (Railo et al., 2021), shaping neural processing before external input is received. Such anticipatory mechanisms have been shown to improve behavioral performance, leading to faster responses and reduced error rates when upcoming events are predictable (Petro et al., 2019), as well as enhance decision confidence (Iemi et al., 2017; Samaha et al., 2017).

However, the mechanisms underlying these behavioral improvements remain debated. One account proposes that anticipation enhances sensory processing (Carrasco, 2011; Iemi et al., 2017; Kok et al., 2012), thereby increasing the quality of evidence available for decision-making and subsequently increasing the rate of evidence accumulation (Kloosterman et al., 2019; Urai et al., 2019; White & Poldrack, 2014). Alternatively, anticipation may bias the decision process itself, for instance by shifting the starting point toward the expected alternative (Bode et al., 2012; Mulder et al., 2012; Yu & Cohen, 2008) or lowering the decision threshold, without directly affecting sensory encoding (Bogacz et al., 2006; Cisek et al., 2009). Distinguishing between these mechanisms is not straightforward at the behavioral level, as both can lead to similar improvements in response times and accuracy. Computational models of decision-making, such as the diffusion decision model, provide a principled framework to dissociate changes in evidence accumulation from changes in decision bias.

Electroencephalography (EEG) has played a key role in the study of anticipatory processes due to its high temporal resolution. In particular, pre-stimulus modulations of event-related potentials (ERPs), such as stimulus-preceding negativities and contingent negative variation (Poulton, 1950; Walter et al., 1964), as well as changes in low-frequency oscillatory activity (Denison et al., 2024; see Tan et al., 2024 for a review), have been linked to anticipatory attention, sensory readiness, and expectation. These effects are often interpreted as reflecting the allocation of attentional resources (Jensen & Mazaheri, 2010), sensory encoding (Hetenyi et al., 2024; Lou et al., 2014), or top-down processing to prepare for upcoming stimuli (Min & Herrmann, 2007). However, much of this literature has focused on averaged signals and univariate contrasts, which reduces trial-by-trial variability. This is particularly limiting for anticipatory processes, which are inherently variable across trials, leaving open the question of whether anticipatory activity carries information about the specific content of upcoming events. Moreover, the unconscious nature of anticipation (Koch & Preuschoff, 2007) complicates the accessibility to the ground-truth knowledge of what was expected, as direct participant questioning might alter neural activity (Trevena & Miller, 2002).

A key unresolved issue is whether pre-stimulus neural activity reflects a generic preparatory state, such as heightened arousal or nonspecific attention, or whether it encodes detailed, content-specific expectations, such as the sensory modality of an anticipated stimulus. Classical ERP components such as the contingent negative variation and modulations of alpha-band activity have often been interpreted as reflecting general preparatory or attentional states, supporting the view of a largely non-specific anticipation signal. More recent studies have reported modality-dependent differences in pre-stimulus EEG activity (Fiorini et al., 2021, 2023; Lucia et al., 2023), suggesting that anticipatory activity may carry more specific information about upcoming events. However, these effects are often subtle and difficult to interpret at the single-trial level. As a result, it remains unclear whether anticipatory states can be reliably distinguished based on their neural signatures prior to stimulus onset, in particular in situations where the modality of the upcoming stimulus is unpredictable. These should however exist, since priors are built even in the absence of statistical regularities in the task (Bévalot & Meyniel, 2024; Yu, 2007; Yu & Cohen, 2008).

Single-trial analyses are pivotal in cognitive neuroscience as they unveil the variability and dynamics of neural activity that are often obscured by averaging across trials (Williams & Linderman, 2021). By examining individual trials, previous studies have unveiled distinct engagement states in perceptual decision-making tasks in mice (Ashwood et al., 2022), as well as the time course of decision strategies in rodents and humans (Roy et al., 2021). Single-trial EEG analyses have been employed to predict memory retrieval success (Nakuci et al., 2023), demonstrating that trial-by-trial fluctuations in neural activity can reveal latent cognitive states, even in the absence of explicit reports, processes that are typically invisible when applying classical signal averaging techniques. Additionally, these analyses facilitate the identification of trial-to-trial behavioral variability (Steinemann et al., 2024), enabling a more nuanced understanding of the neural correlates underlying cognitive functions (Nunez et al., 2017; Ratcliff et al., 2009). As anticipation is believed to be a major source of variability in decision-making (Summerfield & de Lange, 2014; Tarasi et al., 2025), its quantification at the single-trial level is key to understanding decision dynamics.

In the present study, we investigated whether anticipatory EEG activity prior to stimulus onset contains modality-specific information that can be decoded at the single-trial level. Participants performed a task in which visual or auditory targets were anticipated either explicitly via cues or implicitly in uncued trials. We first examined behavioral indices of anticipation, including response times and error rates, and characterized pre-stimulus ERP activity by testing for pre-stimulus cue-related modulations in voltage amplitude using a TFCE-based cluster permutation analysis to identify spatiotemporal regions showing significant anticipatory modulation. We then applied multivariate decoding techniques to assess whether the expected sensory modality could be predicted from pre-stimulus EEG on individual trials. Finally, we tested whether neural decoding generalized to uncued trials and linked anticipatory neural states to behavioral performance using diffusion decision modeling. Together, these analyses provide converging evidence that anticipatory neural activity is both modality-specific and behaviorally relevant.

## Methods

### Participants

42 participants (24 males, 18 females; 5 left-handed, 37 right-handed; aged 20 − 64, mean: 30.43 ± 10.78) with normal or corrected-to-normal hearing and vision took part in this study. The experiment was approved by the Comité d’Ethique de la Recherche Paris-Saclay, under the application number 321. Each participant was informed about the purpose of the study and signed informed consent forms upon participation. All participants fully completed the experiment.

### Experimental design

The experiment consisted of two parts, denominated “uncued” and “cued” phases thereafter, which the participants completed within the same recording session. During the uncued phase, participants were presented randomly at each trial with either the sketch of a face (“face” trial) or a sound (“sound” trial). They were instructed to respond as fast as possible to each stimulus by pressing with their dominant hand on the right arrow of a keyboard for “face” stimuli and on the left arrow for “sound” stimuli. Using the same response modality for both stimuli (click of the dominant hand) ensures that anticipatory effects are not contaminated by motor preparation components. Each trial started with the apparition of a red cross in the middle of the screen, indicating a 1.5 *s* rest period to the participant. The cross then became white to instruct participants to start focusing on the task and avoid parasitic movements that can blur EEG signals (blinking, jaw and head movements in particular). This baseline period lasted for 1.5 − 3 *s*, with its duration varying randomly across trials. After that, a square visual noise clip appeared for 0.9 *s* in the middle of the screen. While the last frame remained on the screen until the end of the trial, the stimulus was displayed for 200 *ms* at the end of the noise clip. The trial terminated upon participant response or after a timeout of 2 *s* after stimulus onset.

Trials in the cued phase were identical to those in the uncued phase, with the addition that a cue appeared for 200 *ms* at the beginning of the noise clip, indicating which stimulus would be displayed. A pictogram of an eye indicated a “face” trial, and an ear icon indicated a “sound” trial. To eliminate the possibility of having created a delayed response instead of sensory anticipation, we set the cues to be inaccurate in 20% of the cases, excluding the ten first trials. We chose this catch probability according to the threshold at which oddball paradigms function, hence ensuring that the cue was still reliable while introducing uncertainty and surprise among subjects (Chavarriaga et al., 2014; Ehrlich & Cheng, 2018; Noah et al., 2020). A summary of the trial sequence for these two phases is given in Figure 1.

**Figure 1.**
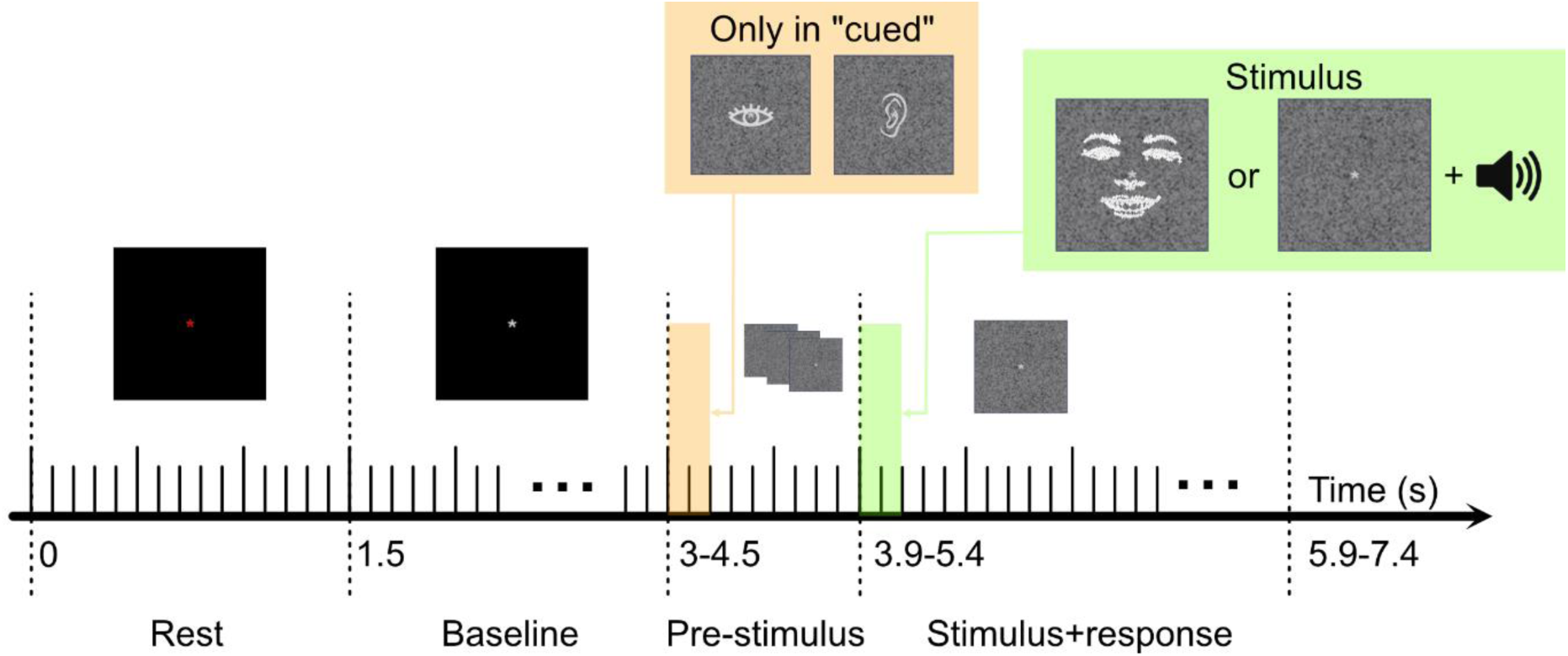
Summary schematic of the sequence of a trial. Each trial starts with a 1. 5 S rest period, followed by a 1. 5 − 3 S baseline period, during which participants are instructed to limit movement and focus on the screen. The pre-stimulus period then lasts for 0. 9 S, a period during which visual noise is presented. In the “cued” condition, the visual cue is presented during the first 200 mS of this period, on top of the visual noise. Then the stimulus, either an image of a face or a sound, is presented for 200 mS in addition to a single frame of visual noise which lasts until the end of the trial. Participants are instructed to report the perceived stimulus as fast and accurately as possible by pressing on the left arrow for a sound and the right arrow for a face stimulus. The trial is terminated upon participant’s response or after a 2 S timeout.

Participants performed 3 uncued and 4 cued blocks of 60 trials each, resulting in 180 uncued trials and 240 cued trials. 20 training trials preceded the uncued phase, and 60 training trials were performed before the cued phase. The training phase preceding the cued phase was longer so that participants had time to learn the impact of the cue, the correct response to each target stimulus, and to avoid making mistakes in catch trials. To prevent any influence of the order of the phases, half of the participants performed the uncued blocks first, while the other group of participants started with the cued blocks. Participants could take a self-paced break between each block. The experiment was performed in a dark room to enhance the sight of the visual stimulus.

### Stimuli

Target stimuli consisted of a sound pulse at 1000 *Hz* for the sound stimulus, and a sketch of a face for the visual stimulus. The sketches were generated by Yang et al. (2020) from the Radboud Faces Database (Langner et al., 2010). Cue stimuli consisted of an open-source drawing of an eye and one of an ear, displayed at the center of a screen over a random dot square cloud. All stimuli lasted for 200 *ms*.

### EEG data acquisition

EEG signals were recorded at 1000 *Hz* using 32 active AgCl electrodes, and the actiCHamp Plus amplifier (Brain Products GmbH, Gilching, Germany). The electrodes were placed according to the 10/20 international system. An electrode located at *FCz* served as the reference electrode upon acquisition.

### Time-domain EEG comparisons

Event-related potentials (ERPs) were analyzed to assess whether anticipatory neural activity could be distinguished in the time domain between anticipation and no anticipation, as well as between visual and auditory anticipation. The pre-stimulus period was defined as the window [−400: 0] *ms* preceding the target stimulus for both stimuli and in both cued and uncued conditions. It can equivalently be defined as the [500: 900] *ms* time window following the onset of the noise clip and cue. 2 comparisons were performed: anticipation vs. no anticipation (defined as cued vs. uncued trials), and visual vs. auditory anticipation (defined in the cued condition as the type of cue presented: eye or ear).

EEG data were low-pass filtered at 35 Hz using a minimum-phase finite impulse response (FIR) filter to focus on event-related activity while preserving temporal precision and hence avoiding contamination by post-stimulus activity. Data were then epoched relative to pre-stimulus period onset. Linear detrending was applied at the epoch level to remove slow drifts, rather than relying on high-pass filtering, in order to avoid introducing distortions in pre-stimulus activity and anticipatory signals. A conservative artifact rejection was performed automatically by excluding epochs with peak-to-peak amplitudes exceeding ±200 µV (Delorme, 2023). To preserve sufficient trial counts while maintaining data quality, an adaptive procedure was used in which electrodes were iteratively removed and interpolated them using spherical spline interpolation, when trial rejection exceeded 20% for a given participant. We ensured that fewer than 4 channels were interpolated for each participant (corresponding to ∼10% of channels).

For the initial investigation of significant differences between the different periods of interest described above, ERP components were computed by averaging trials within each category (cued trials, uncued trials, face anticipation, and sound anticipation).

### Classification pipeline

To assess whether anticipatory neural activity contained information about the expected sensory modality at the single-trial level, we performed a multivariate classification analysis on pre-stimulus EEG data. Spatial filters were computed using the XDawn algorithm (Rivet et al., 2009) for each class using the train set, selecting 5 components which went through minimum-maximum scaling. Logistic regression with L1-regularization was then applied to predict the class of each trial. This classification method is based on building spatial filters to split data into categories. Since we expect that visual and auditory anticipation utilize different networks in the brain, leveraging the spatial structure of data is relevant. Because anticipatory EEG activity has been reported across multiple low-frequency ranges and no strong a priori frequency band could be specified, we performed an exploratory grid search over several canonical frequency bands (delta (1-4Hz), theta (4-8Hz), and alpha (8-13Hz) ranges), as well as a 35Hz low-pass filter, using only cued trials. Cued trials were first partitioned into separate training and test sets using a stratified shuffle split. The test set consisted of 25% of the trials for each participant (i.e. 60 trials). Frequency-band selection was performed exclusively within the training set using stratified 10-fold cross-validation. This analysis was used to characterize the spectral range carrying anticipatory information and to inform the selection of a single frequency band for subsequent generalization analyses. To determine a single frequency band for subsequent analyses, we averaged cross-validation accuracy across participants for each band and selected the band yielding the highest mean validation performance. Importantly, the final decoding models were trained on the full training set for each participant and evaluated on the held-out test set.

#### Classifying pre-stimulus activity of uncued trials

The afore-described classification pipeline is based on supervised learning, i.e. it requires the knowledge of a ground-truth. In this study, it requires a knowledge of the type (visual or auditory) of anticipation. However, the ground-truth label of anticipation is unknown on uncued trials. To circumvent this issue, if we assume the stationarity of the anticipation phenomenon across phases of the task, the brain activity patterns of anticipation over uncued trials should be similar to those found on the cued trials. Therefore, classifiers trained on cued data can be applied on uncued data of the same participant. We hence re-trained the classifiers obtained on all the trials of the cued phase of the experiment for each participant and predicted the corresponding anticipation class using the thus trained classifier on uncued trials. We treat the classifier’s single-trial predictions as an index of putative anticipatory priors while explicitly noting that this index does not by itself prove a belief-like computation without additional control analyses.

### Linking brain and behavior

One of the main assumptions regarding the behavioral effects of anticipation is that correct expectations lead to reduced response times and error rates. Several mechanisms have been proposed to account for these effects. On some accounts, anticipation increases sensory sensitivity, thereby enhancing the quality of evidence accumulation during decision formation (Iemi et al., 2017; Kloosterman et al., 2019; Samaha et al., 2017; Urai et al., 2019). Other accounts instead suggest that anticipation biases the decision process by shifting the starting point toward the expected alternative (or, equivalently, lowering decision threshold), without directly affecting sensory evidence (Bode et al., 2012; Mulder et al., 2012). Distinguishing between these mechanisms requires a computational framework that can dissociate changes in evidence quality from changes in decision bias. If the classification algorithm was indeed trained to distinguish the type of anticipation, we should observe that correct anticipations are more represented among shorter reaction times relative to long ones. Conversely, incorrect anticipations should result in longer reaction times.

To analyze the effects of anticipation on behavior, we fitted diffusion decision models (DDMs) (Ratcliff, 1978; Ratcliff & McKoon, 2008; Ratcliff & Tuerlinckx, 2002) to behavioral data. According to DDMs, sensory evidence is accumulated linearly from a starting point *z*_*r*_ until reaching a fixed decision boundary a, at which time a decision is made.

The DDM is defined by the equation:

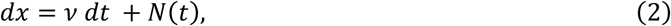

where the decision variable *x* varies by *dx* in infinitesimal time *dt*. The decision state can be viewed as a particle subject to Brownian motion with drift *v*, Gaussian white noise term *N*(*t*). Additionally, a non-decision time *T*_0_is fitted to account for the biological delay of sensory encoding and motor preparation explaining the difference between the decision time and the observed response time.

Our DDM analysis aimed at identifying whether changes in anticipation resulted in changes in the accumulation rate *v*, or rather in a bias in the starting point *z*_*r*_. More specifically, correct anticipation resulting in faster and more accurate decisions than incorrect anticipation, one could expect that correct anticipation either increases the rate of evidence accumulation, or that the decision is initially biased towards the correct decision.

On cued data, we hypothesized that correct anticipations should result in larger drifts or larger starting points than incorrect anticipations. To test this hypothesis, we fitted a DDM for each participant, fixing the boundary *a* = 1 and non-decision time *T*_0_ = 0.3 *s* for all participants. This was necessary in order to compare model parameters across participants, since these parameters are interdependent with the drift and the starting point. For each participant, separate drift terms and starting points were fitted depending on whether ground-truth anticipation was correct or incorrect. The software pyDDM was used for this analysis. We then performed paired-sample -tests to assess whether the drifts and starting points significantly depended on the correctness of anticipation.

The same DDM fitting was performed on uncued data, using this time the anticipation class predicted by the classifier to assess the correctness of anticipation. This serves as a validation step to the classification: if we observe the same trends as in the cued dataset, this would indicate that the classifier captures anticipatory neural states that have functional consequences for decision formation, rather than reflecting task-specific or stimulus-locked confounds.

### Statistical analyses

#### Behavioral effects

We first tested the effects of the presence of the cue and the stimulus on the response time and accuracy at the group level using a repeated measures ANOVA procedure, with “Stimulus type” (auditory, visual) and “Condition” (cued, uncued) as within-subject factors. We further tested the effect of congruent and incongruent cues and stimuli on the response time and accuracy on the cued condition only, using repeated measures ANOVA again, with “Stimulus type” and “Cue congruency” as within-subject factors. Note that the tests were done separately in each condition due to the unbalanced number of trials.

#### Significance of time-domain differences

Group-level statistical significance of anticipatory effects in the ERP time series was assessed using threshold-free cluster enhancement (TFCE, Smith & Nichols (2009)) on ERP differences combined with non-parametric permutation testing, as implemented in MNE-Python (Maris & Oostenveld, 2007). This approach evaluates effects that are both strong and extended across time and electrodes while controlling for multiple comparisons. TFCE avoids the need to define an arbitrary cluster-forming threshold by integrating information across a range of thresholds. For each comparison, one-sample t-statistics were computed at each channel and time point across participants. Condition labels were randomly permuted to generate a null distribution of TFCE values, and for each permutation the maximum TFCE value across all spatio-temporal points was retained. Observed TFCE values were compared against this null distribution to determine statistical significance. A total of 1024 permutations were performed, providing a stable estimate of the null distribution while remaining computationally efficient. For visualization, spatio-temporal clusters belonging to TFCE effects with a corrected p-value below 0.05 were marked as significant.

#### Significance of classification performance

We compute the classification accuracy on the test set for each participant on cued trials. The classification scores are compared to chance level that we obtained through permutation testing over 1000 iterations. At each iteration, the labels are shuffled, and the classifier is re-run to compute the classification accuracy over the randomly labeled data. The null-hypothesis distribution of the chance level classification performance is thus obtained for each participant, and the classification performance is compared against this distribution. The *p*-value of the classification accuracy against the null-hypothesis distribution is computed for each participant as 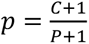, with *C* the number of permutations for which the accuracy is superior or equal to the test accuracy, and *P* = 1000 the number of permutations. The accuracy indeed needs to lie above the individual chance level to consider that the classification succeeded.

## Results

We analyzed EEG recordings from 42 participants who performed a sensory categorization task. At each trial, participants decided whether the randomly presented was a sound or a visual stimulus consisting of a drawing of a face. Each trial was preceded by a pre-stimulus period lasting 0.9 seconds. On some trials, a cue was presented at the start of this pre-stimulus period, indicating with 80% confidence the class of the upcoming stimulus. In such cases, the pre-stimulus period was referred to as the *anticipation* period.

Our analyses aimed to characterize brain activation patterns associated with specific stimulus anticipation, both at the group level and at the single-trial level.

### Cued anticipation yields faster responses

The descriptive statistics of response times and accuracies across conditions and stimulus types are summarized in Figure 2 (A and B). We first tested the effects of stimulus type and condition (i.e. presence or absence of an informative cue) on the response times and accuracy at the group level using ANOVA. We observed a significant effect of stimulus type (*RT*_*auditory*_ = 526*ms*, *RT*_*visual*_ = 462*ms*, *F*_{1,41}_ = 119.783, *p* < 0.001) and condition (*RT*_*cued*_ = 470*ms*, *RT*_*uncued*_ = 526*ms*, *F*_{1,41}_ = 70.304, *p* < 0.001) on response times. Additionally, we identified an interaction effect between stimulus type and condition (*F*_{1,41}_ = 30.024, *p* < 0.001), with responses of cued visual trials faster than any other trial type (*p* < 0.001). Uncued visual trials were significantly faster than uncued auditory trials (*t* = 12.236, *p* < 0.001), but not faster than cued auditory trials (*t* = 1.058, *p* = 0.293).

**Figure 2.**
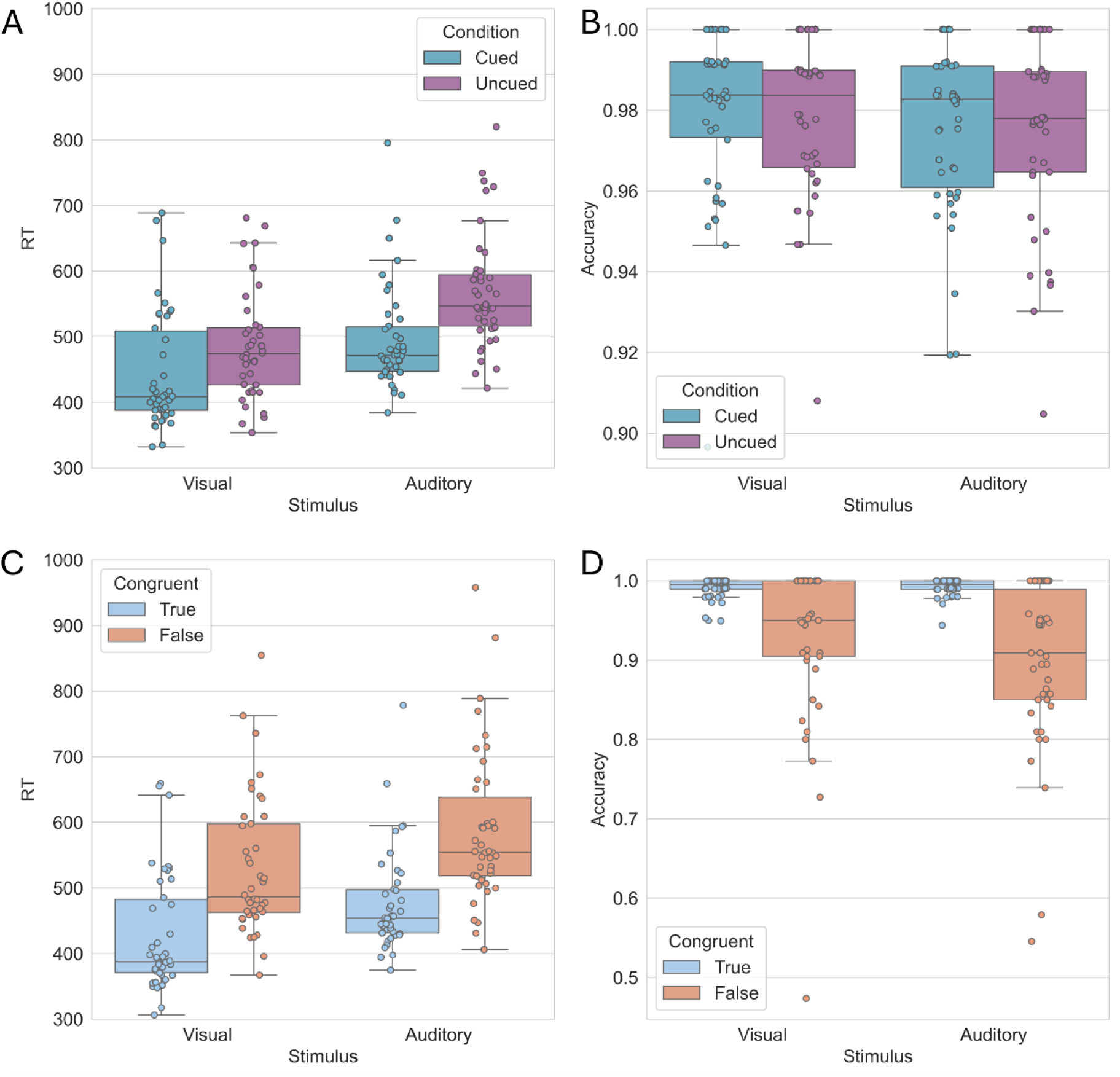
Mean response times in mS (A, C) and accuracy (B, D) for all participants for each stimulus type, depending on (A, B) the cued/uncued condition and (C, D) whether the cue was congruent or not in cued trials.

In contrast, the accuracy was not affected by either stimulus type or the presence of the cue (see Figure 2B). Furthermore, we observed no interaction effect between the two factors on accuracy. These results show that the presence of the cue effectively reduced response times (see also Figure 2A).

Next, we examined the effect of congruent and incongruent cue, as well as stimulus type, on the response times and mean accuracy within the cued condition using ANOVA (see descriptive statistics in Figure 2C and 3D). Incongruent trials had a significant effect on both response time (*RT*_*congruent*_ = 452*ms*, *RT*_*incongruent*_ = 558*ms*, *F*_{1,41}_ = 131.890, *p* < 0.001) and accuracy (*Acc*_*congruent*_ = 99.1%, *Acc*_*incongruent*_ = 91.1%, *F*_{1,41}_ = 31.140, *p* < 0.001). As previously, stimulus type influenced response times (*RT*_*auditory*_ = 496*ms*, *RT*_*visual*_ = 445*ms*, *F*_{1,41}_ = 67.009, *p* < 0.001, see also Figure 2C) but did not affect accuracy (Figure 2D). No interaction effect was found on either response times or accuracy. These results demonstrate that participants considered the cue to be informative.

**Figure 3:**
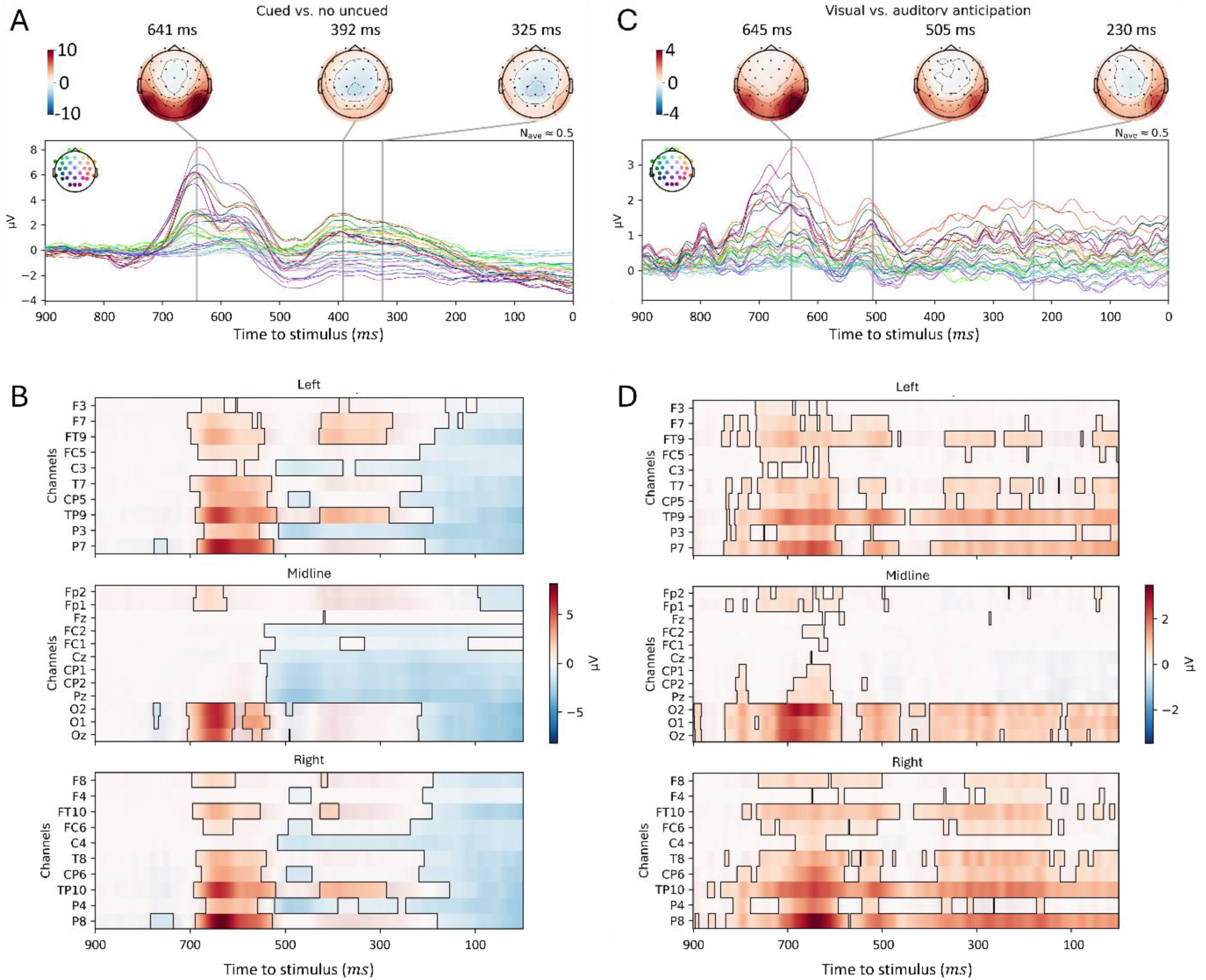
ERP difference traces of pre-stimulus activity. (A) cued vs. uncued trials. The topographies and traces represent cued activity minus uncued activity. (B) spatio-temporal difference of activity between cued and uncued trials. The outlined clusters represent the significant results of TFCE analysis, with p < 0. 05 according to permutation testing. (C) cued trials are split on whether the cue was the drawing of an eye (“visual” anticipation) or an ear (“auditory” anticipation). The topographies and tracesrepresent visual minus auditory anticipatory activity. (D) spatio-temporal difference of activity between visual and auditory anticipatory activity. The outlined clusters represent the results of TFCE analysis with p < 0. 05 according to permutation testing. On the cluster analyses (B and D), electrodes are grouped whether they are on the left (top), midline (middle) or right (bottom) of the scalp, with the midline encompassing ***z***, 1 and **2** electrodes.

Together, these results support the emergence of anticipatory effects following the presentation of an informative but partially unreliable cue, thereby validating the experimental paradigm implemented in this study.

### Pre-stimulus EEG activity reflects contents beyond temporal anticipation

We first sought to demonstrate that the anticipation activity goes beyond temporal expectation of the coming event. To this end, we compared brain activity during the anticipation period of cued and uncued trials at the group level.

ERP differences of the pre-stimulus period (Figure 3A) reveal two notable activation differences. The first one, occurring at approximately 650ms to stimulus, corresponds to the visual-evoked potential due to the presentation of the cue. This activity reflects sensory processing of the cue, making it irrelevant for the subsequent analyses. The second time window, the last 400ms before stimulus onset, shows a ramping of activity, which is typically observed during temporal expectation. Since this signal differs from that observed in uncued trials - where perceptual anticipation is minimal and temporal expectation prevails - it suggests that the content of anticipation, specifically the expected sensory modality, shapes the associated brain activity.

Figure 3B also highlights the areas involved in anticipation by illustrating the difference in evoked activity during the pre-stimulus period between the cued and uncued phases, i.e. with and without an informative cue. We observe a general ramping of EEG activity, and a more negative difference in the central areas. Later activity is dominated by occipital and central activation. The highlighted parts of that figure represent significant cluster, according to the TFCE+permutation testing method.

Next, we investigated whether there existed differences within cued trials between visual and auditory anticipation, that is, between trials cued with an image of an eye and those cued with an image of an ear. Differences in ERP (Figure 3C) show three periods of interest: one around 650ms before stimulus onset, which could again represent the P300 component of cue presentation, a secondary peak at 505ms pre-stimulus, and an occipital and temporal ramping in the last 400ms of the pre-stimulus window. Our TFCE analysis (Figure 3D) reveals significant differences in the spatio-temporal pre-stimulus activity between these two types of trials.

Notably, we observe that the amplitude and spatialization of the difference in ramping in the last 400ms is broader across conditions than within the cued condition, which suggests that this activity relates to the strength of stimulus anticipation. Indeed, participants expect a given stimulus more strongly if it is cued than if uncued.

We additionally investigated the relationship between electrode-wise activity in the pre-stimulus and post-stimulus periods to assess whether anticipatory effects spatially correspond to stimulus-evoked responses. For each participant, cued trials were separated according to the presented cue and averaged across trials to obtain condition-specific ERPs. We then computed, for each electrode, the difference in ERP amplitude between conditions (auditory vs. visual). Pre-stimulus activity was defined as the average signal in the −400 to 0 ms window relative to stimulus onset, and post-stimulus activity as the signal in the 0 to 400 ms window. The absolute values of these condition differences were then computed and averaged across time within each window. Finally, for each participant, we computed the correlation across electrodes between pre-stimulus and post-stimulus activity, yielding a measure of the spatial correspondence between anticipatory and evoked effects.

We found that electrodes showing stronger anticipatory differences between conditions also tended to exhibit stronger stimulus-evoked differences (Figure 4A–B), with posterior electrodes displaying greater cue-related effects. Regression slopes were consistently positive, indicating that electrodes with larger anticipatory effects also showed larger stimulus-evoked responses. This relationship was robust at the group level, with slopes significantly greater than zero across participants (Figure 4C, mean slope: 1.98, 95% CI [0.71, 3.44], one-sample *t*-test against zero: *t*_41_ = 16.12, *p* < 0.001) and the model explaining a substantial proportion of variance (Figure 4D, mean *R*^2^ = 0.71, 95% CI: [0.34, 0.93]).

**Figure 4:**
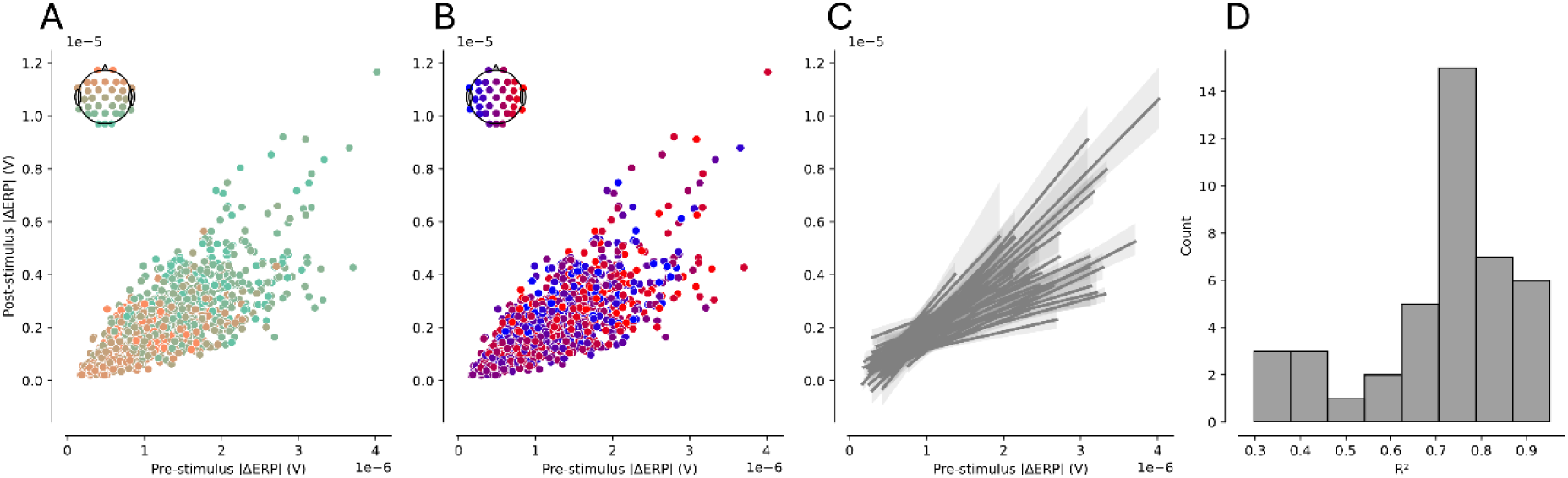
Spatial correspondence between pre-stimulus and post-stimulus ERP differences between eye-cued and ear-cued trials, across electrodes. (A-B) Electrode-wise relationship between pre-stimulus and post-stimulus ERP differences. Each dot represents one electrode for one participant, and is color-coded according to position along the rostral-caudal axis (A) and medial-lateral axis (B). (C) Linear regression computed separately for each participant across electrodes, showing consistent positive slopes. Shaded areas indicate 95% confidence intervals. (D) Distribution of explained variance (*R*^2^coefficient) across participants.

This result supports the hypothesis of anticipation as a pre-activation of relevant processing areas.

### Pre-stimulus activity can successfully be classified according to anticipation content

We next classified pre-stimulus activity of cued trials according to the type of cue presented. Importantly, we considered only the 400ms preceding stimulus onset (i.e. the period [500ms – 900ms] after cue onset) to not classify the P300 component related to the detection of the cue, and as guided by our ERP analysis. A grid-search on the train set indicated that filtering in the 4-8Hz band yield higher accuracy across all participants, so trials in the test set were filtered in that frequency band.

The average classification accuracy on the test set was 0.66 ± 0.08 (mean±standard deviation), with data from 31/42 participants classified above their empirical chance level, computed by means of permutation testing (Figure 5), leaving data from 11/42 (i.e. 26%) classified at chance level.

**Figure 5:**
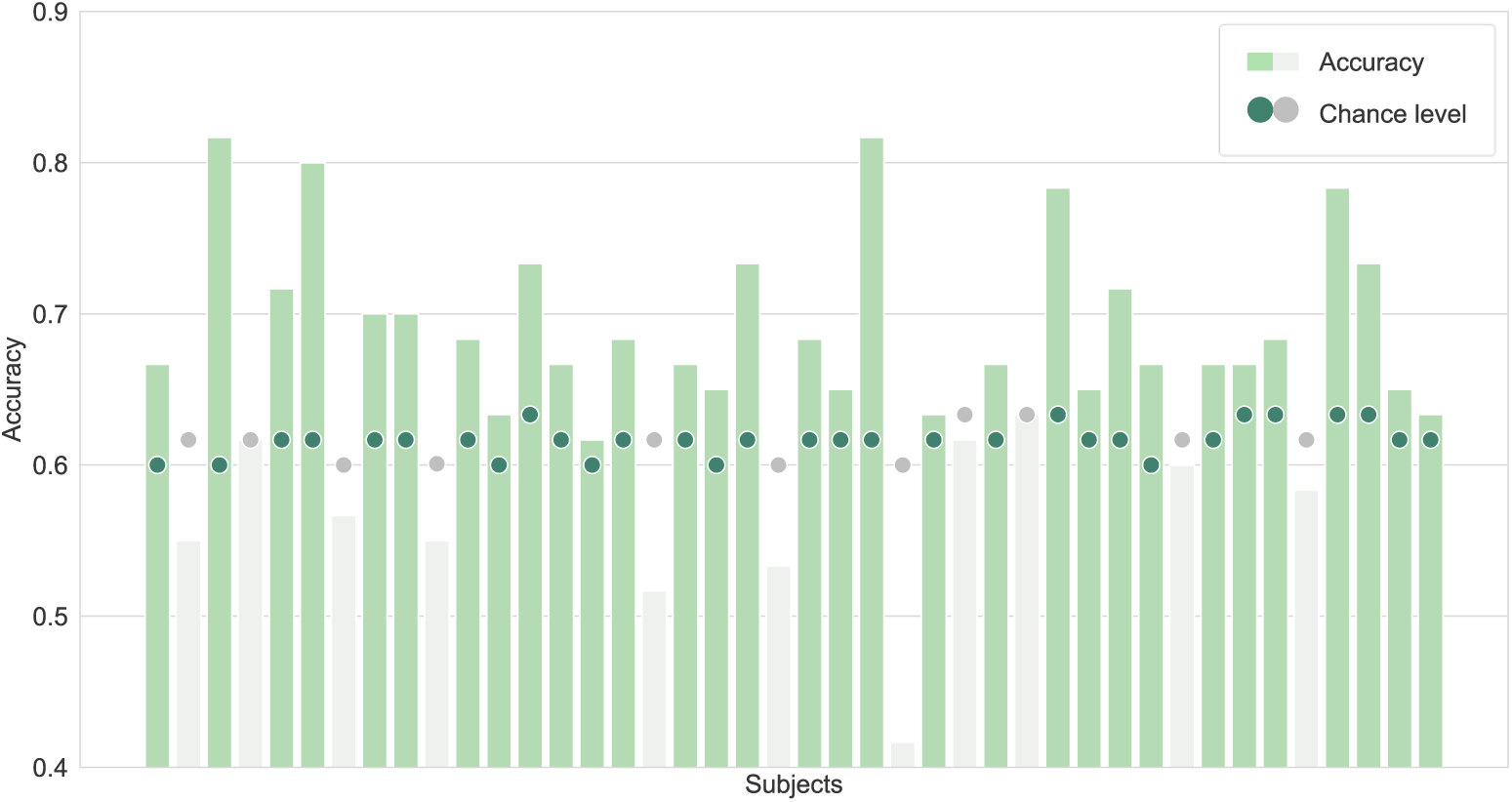
Classification accuracy obtained from XDawn+Logistic Regression classification of the 4 − 8*Hz*-filtered pre-stimulus activity of cued trials (”accuracy”, bars), compared to the empirical chance level (”chance level”, dots) computed by permutation testing (1000 permutations). Subjects for which the classification accuracy is above chance level are highlighted in green.

We investigated whether the difference in classification accuracy could be related to inter-subject differences in task design or demographics. We found that, out of the 11 participants whose anticipation could not be decoded by the classifier, 6 saw the cued blocks first and 5 the uncued blocks first, 1 was left-handed (i.e. we obtained chance-level classification accuracy for 20% of left-handed participants), 4 were female and 7 were male (i.e. we obtained chance-level classification accuracy for 22% of females and 29% of males).

To test statistically the dependence of classification accuracy on inter-subject differences, we performed two-sample *t*-tests, revealing no difference in classification accuracy related to block order (*t* = 0.941, *p* = 0.352, Figure 6A) or sex (*t* = 0.697, *p* = 0.490, Figure 6B). To test whether classification accuracy depended on handedness, we compared the mean accuracy of left-handed participants to a null distribution obtained by randomly sampling 5 accuracies (matching the number of left-handed participants) from the full sample and averaging them. This procedure was repeated 1000 times to generate the null distribution. The observed mean accuracy of left-handed participants did not differ significantly from this distribution (*p* = 0.321, Figure 6C). Finally, a regression analysis outlined no significant relationship between classification accuracy and age (*r* = 0.071, *p* = 0.655, Figure 6D). These results are summarized in Figure 6.

**Figure 6:**
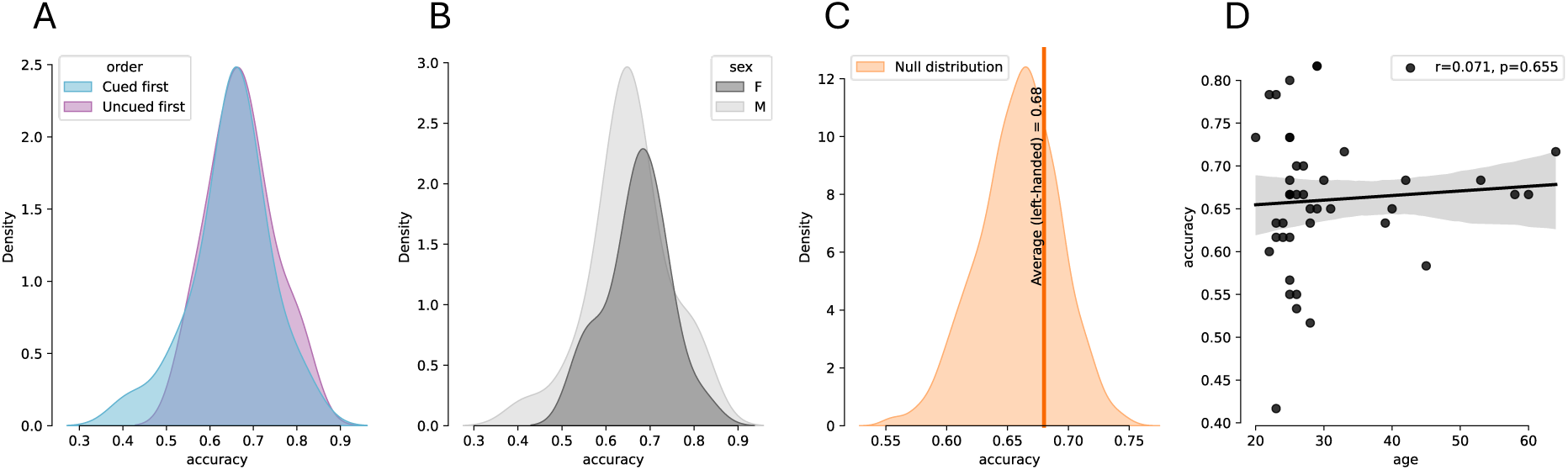
Inter-subject differences in classification accuracy, depending on block presentation order (A), sex (B), handedness (C) and age (D). We find that none of these factors relate to classification accuracy.

The same classifiers were retrained on all the cued trials per participant and applied to the uncued trials to obtain an index of putative anticipation. In the next section, we explain how the computed labels were validated using behavioral analyses.

### Anticipation and diffusion-decision models

We fitted the Diffusion-Decision Model (DDM) to individual behavioral data in the cued condition to examine the mechanism by which response times and response accuracy improve when the upcoming stimulus is correctly anticipated. For this analysis, the boundary *a* = 1 and non-decision time *T*_0_ = 0.3*s* were fixed for all participants, while the drift *v* and starting point *z*_*r*_ were fitted individually for each participant, depending on whether the cue matched the stimulus or not. This resulted in two drifts and two starting points per participant.

We first tested whether the drift or the starting point varied significantly based on the correctness of anticipation. The starting point was significantly closer to the correct decision boundary when the stimulus was correctly anticipated (Shapiro-Wilk test: *W* = 0.862, *p* < 0.001, Wilcoxon signed rank test: *W* = 879, *z* = 5.345, *p* < 0.001, matched rank biserial correlation (effect size): *r*_*rb*_ = 0.947). No significant difference was observed between drift parameters.

#### Assessing Classification Performance on Uncued Data

To evaluate classification performance on uncued data, we repeated the subjective fitting procedure using uncued trials. We computed the difference between the parameters (drift rate and starting point) of correctly and incorrectly anticipated trials, using the predicted anticipation class to split the trials. Figure 7 shows the parameter differences.

**Figure 7:**
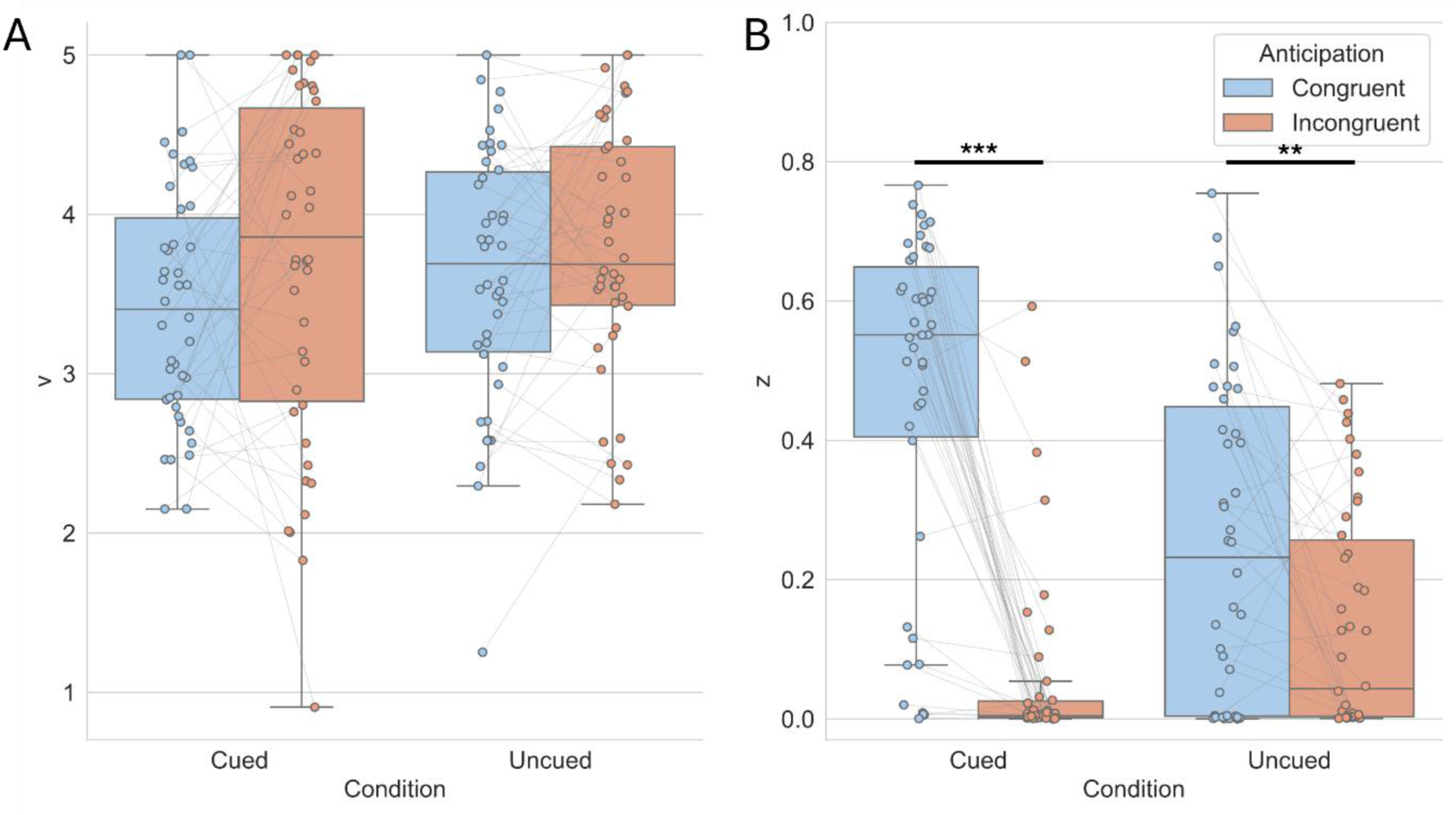
DDM parameter differences between congruent (in blue in both plots) and incongruent (in orange in both plots) anticipation. (A) differences in drift term *v* are not significant between anticipation types, whereas we observed notably higher bias ***z*** (B), i.e. bias toward the correct decision boundary, in correctly compared to incorrectly anticipated trials. ***:*p* < 0. 001, **:*p* < 0. 005.

We observed once again a significant difference in starting points, with the starting point of correctly-anticipated trials closer to the decision boundary than incorrectly-anticipated (Shapiro-Wilk test: *W* = 0.934, *p* = 0.017, Wilcoxon signed rank test: *W* = 682, *z* = 2.882, *p* = 0.003, matched rank biserial correlation (effect size): *r*_*rb*_ = 0.511). No significant difference was observed between drift parameters. These results being consistent with the observations made on cued trials supports that the classifier picked relevant differences in pre-stimulus activity in uncued trials. We note that the parameter distribution in biases is wider in the uncued condition, which we can attribute mainly to the classification, which is approximate. Nevertheless, the consistency of results between the cued and uncued condition is an indicator that the classification caught relevant variations in the pre-stimulus period.

## Discussion

The present study investigates whether anticipatory neural activity prior to stimulus onset reflects modality-specific expectations and whether such activity is functionally relevant for behavior. Using a combination of ERP analyses, multivariate decoding, and computational modeling, we show that pre-stimulus EEG activity differs between visual and auditory anticipation, can be decoded at the single-trial level, and predicts behavioral performance even in the absence of explicit cues. These findings support the view that anticipation is not merely a nonspecific preparatory state, but instead involves content-specific neural representations – here at the level of sensory modality - that shape subsequent decision processes.

At the neural level, ERP analyses revealed significant differences in pre-stimulus activity between anticipation conditions. Importantly, these effects were observed prior to target onset, ruling out contamination from stimulus-evoked responses. Our cluster analyses reveal that anticipatory activity is distributed, which is consistent with findings in the mouse that prior information is encoded in various brain areas (Findling et al., 2025). Importantly, we observe significant differences not only between cued and uncued trials, but also within cued trials depending on what cue has been presented. This finding extends earlier accounts that associated ERP pre-stimulus activity with non-specific temporal expectation (Walter et al., 1964), temporal estimation (Kononowicz & Penney, 2016). Rather than reflecting a purely temporal readiness signal, our findings indicate that pre-stimulus ERP activity can carry information about the content of upcoming sensory events, specifically their sensory modality. This interpretation aligns with more recent works distinguishing pre-stimulus activity related to temporal expectation and temporal attention (Denison et al., 2024), and that anticipatory theta activity relate to causal predictions in infants (Begus & Bonawitz, 2024). In that line, our ERP analysis suggests that anticipatory activity links to predictions of the upcoming event, which are stronger when reliable cueing is presented and attenuated in uncued trials.

Crucially, pre-stimulus activity was successfully decoded for 31 out of 42 participants displaying above-chance single-trial classification performance in cued trials. Other works demonstrated to potential to differentiate anticipation at the single-trial level in Go/NoGo tasks, specifically distinguishing trials requiring motor responses from those in which responses are withheld (Chavarriaga et al., 2012; Garipelli et al., 2009, 2011; Khaliliardali et al., 2012, 2015). Notably, Chavarriaga et al. (2012) eliminated the influence of motor preparation by including tasks without motor output, arguing that the differences are ascribable to attentional shifts following errors. Our findings extend these results by demonstrating that the pre-stimulus activity reflects stimulus-specific anticipation, opening new insights into the complexity of anticipatory processes. Other works, in particular those studying language, have found that pre-stimulus activity is modulated by word predictability (Kölbl et al., 2025; León-Cabrera et al., 2024). In our work, while cued trials indicate an 80% probability of having the corresponding stimulus, there is a 50% chance of either stimulus occurring at any time in uncued trials. The apparent distinguishability between visual and auditory expectation even in uncued trials aligns with previous accounts that participants estimate probabilities even in the absence of statistical regularities (Yu, 2007). Arguably, participants in our work may have displayed stronger differences if they were not instructed about the task (Bévalot & Meyniel, 2024). However, the training of classifiers in our work relies on the existence of cued blocks, which have to be displayed at the beginning or the end of the experiment to counterbalance learning effects. Future works, notably using unsupervised learning techniques, could test this hypothesis.

This finding goes beyond traditional ERP analyses by showing that anticipatory states contain sufficient information to distinguish between visual and auditory expectations at the level of individual trials. Importantly, Fiorini et al. (2023) have compared conditions where the modality of the cues matched the modality of the stimulus. In one of the conditions, the cues were mixed such that the second stimulus was unpredictable. They show that there are differences in the aP and vN components depended on the conditions in which they were presented. However, their observations cannot completely discard the influence of sensory processing of the cue in the ERP traces. While we observed a similar ramping of activity (which was also observed in other tasks and characterized as temporal attention (Denison et al., 2024)), we managed to classify them at the single-trial granularity, even in the absence of any informative cue. Barik et al. (2019) have classified single-trial alpha-band activity in face pareidolia, showing that the detection of faces in the absence of evidence relates to lateralized alpha-band activity. Their approach however does not allow to distinguish between motor processes (they indeed observe a lateralization of alpha-band activity, consistent with motor mu-band lateralization) and anticipation. By using the same response modality for both stimuli (click of the dominant hand), our results cannot be explained by motor preparation components.

The generalization of decoding performance to uncued trials provides further insight into the nature of anticipatory processes. Even in the absence of explicit cues, classifier predictions remained were behaviorally meaningful, suggesting that participants formed endogenous expectations based on perceived task structure. This result argues against a purely cue-driven interpretation of anticipatory signals and instead supports the idea that the brain continuously generates predictions about upcoming sensory events, even when such predictions are implicit. By linking neural decoding to diffusion decision model parameters, our findings offer a mechanistic account of how anticipation influences behavior. Correct anticipatory states were associated with higher bias, indicating more efficient accumulation of sensory evidence. Previous works have also established a relation between starting point and previous trials (Bode et al., 2012; Grosjean et al., 2001), while others have discovered higher drift rates with higher expectations (Urai et al., 2019) (although see covariation of drift and bias in Hoxha et al. (2023)). We note that these studies were only performed in the visual modality, and that parallel networks may be at play in our bimodal task, hence the absence of significant difference between drift terms.

Decoding performance was highest when signals were filtered in the theta band (4–8 Hz), suggesting that anticipatory information is preferentially carried by low-frequency neural activity. Importantly, our analysis was conducted in the time domain, and therefore does not directly speak to ongoing oscillatory activity per se, but rather to slow fluctuations in pre-stimulus neural signals. This finding is nevertheless consistent with a large body of literature linking theta oscillations to top-down control (Cavanagh & Frank, 2014; Sauseng et al., 2010) and predictive processes (Arnal & Giraud, 2012; Engel et al., 2001). Theta-band activity has been associated with the coordination of distributed neural networks during expectation and attention (Galas et al., 2025; Michel et al., 2022; Von Stein & Sarnthein, 2000), as well as with the maintenance of task-relevant information over time (Hsieh & Ranganath, 2014; Lisman & Jensen, 2013). In particular, low-frequency oscillations are thought to support long-range communication between frontal and sensory regions (Arnal et al., 2015; Cohen et al., 2008; Fries, 2005), making them well-suited for conveying anticipatory signals prior to stimulus onset (Cohen & van Gaal, 2013). In the context of temporal and sensory expectation, theta activity has also been implicated in aligning neural excitability with expected events, thereby facilitating upcoming processing (Cravo et al., 2013; Schroeder & Lakatos, 2009). In line with this interpretation, our regression analysis revealed that electrodes exhibiting stronger anticipatory differences also showed stronger stimulus-evoked differences, suggesting a spatial correspondence between pre-stimulus and post-stimulus effects. This result is consistent with the idea that anticipatory low-frequency activity may bias subsequent sensory processing by modulating the excitability of task-relevant neural populations prior to stimulus onset, a result that has already been demonstrated with oscillatory activity in temporal expectation (Barne et al., 2021).

One alternative interpretation of the decoded pre-stimulus signal is that it reflects sustained maintenance of the cue representation rather than a predictive sensory template per se. In this view, the classifier would discriminate trials based on the continued representation of the visual or auditory cue, without necessarily indexing preparatory activation of modality-specific sensory systems. Although this account would require maintenance of cue-related information for one second, such sustained representations are not implausible, particularly in task contexts encouraging active retention (Jung et al., 2025; Li et al., 2025). Importantly, the present data are formally compatible with both interpretations. However, several aspects of our findings argue in favor of a predictive account. First, decoding generalized to uncued trials, where no explicit cue representation was available to maintain, suggesting that similar neural states can arise in the absence of a physical cue. Second, the observed modulation of decision parameters indicates functional consequences of the decoded state, consistent with a preparatory role rather than passive maintenance. Future work with a total absence of cue will be necessary to adjudicate between sustained cue maintenance and genuinely anticipatory sensory templates more directly.

A related assumption underlying our generalization analysis is that the neural patterns decoded in cued trials reflect the same type of anticipatory states that occur in uncued trials. Although cross-condition generalization suggests partial overlap between these states, it does not formally demonstrate their equivalence. Successful generalization indicates that the classifier trained on cued trials can detect informative structure in uncued data, but this could arise from shared components embedded within broader, potentially distinct neural configurations. In other words, uncued anticipatory states may recruit similar low-frequency spatial patterns while differing in their strength, stability, or additional contextual components. Future work using representational similarity analysis or cross-temporal generalization approaches could more directly test whether cued and uncued anticipation rely on identical neural codes or instead on partially overlapping but distinct anticipatory mechanisms.

Why do we anticipate, even when that leads to longer reaction times upon incorrect anticipation? Indeed, in ecological environments, it is more likely than not that anticipation is incorrect, as cues are generally absent. Previous works in have suggested that biasing decisions given local patterns is a robust strategy in changing environments, in a Bayesian framework (Yu & Cohen, 2008) and in a reinforcement-learning framework (Hoxha et al., 2025). Our findings provide a neural counterpart to these theoretical perspectives. We show that anticipatory states are expressed as modality-specific patterns of pre-stimulus EEG activity that influence subsequent behavior, even in the absence of explicit cues. Crucially, these anticipatory signals were associated with both benefits and costs: correct anticipations were linked to faster responses and reduced error rates, whereas incorrect anticipations incurred performance penalties. This trade-off suggests that anticipatory neural states do not merely reflect strategic guessing, but rather a proactive biasing of sensory and decision-related processes that persists across trial contexts.

The present results contribute to a growing body of evidence supporting the role of predictive processes in perception and decision-making. By demonstrating that pre-stimulus EEG activity encodes modality-specific expectations that generalize beyond explicit cues and influence decision dynamics, this study highlights the functional significance of anticipatory neural states. More broadly, our findings underscore the importance of studying neural activity prior to stimulus onset to understand how the brain prepares for and ultimately shapes perception and action. Finally, by identifying robust neural features of anticipatory states in cued and uncued contexts, this work provides a principled starting point for future studies to characterize anticipation in the absence of explicit cues, for instance by leveraging unsupervised or data-driven clustering approaches to dissociate distinct anticipatory states.

## Code and data availability

The raw data presented in this study (Hoxha et al., 2026) is available at https://zenodo.org/records/19595833 and the processing code is available at https://github.com/IsabHox/AnticipatoryModality.

